# Habitat loss and fragmentation increase realized predator-prey body size ratios

**DOI:** 10.1101/461376

**Authors:** Jasmijn Hillaert, Martijn L. Vandegehuchte, Thomas Hovestadt, Dries Bonte

## Abstract

In the absence of predators, habitat fragmentation favors large body sizes in primary consumers with informed movement due to their high gap-crossing ability. However, the body size of primary consumers is not only shaped by such bottom-up effects, but also by top-down effects as predators prefer prey of a certain size. Therefore, higher trophic levels should be taken into consideration when studying the effect of habitat loss and fragmentation on size distributions of herbivores.

We built a model to study the effect of habitat loss and fragmentation within a simple food web consisting of (i) a basal resource that is consumed by (ii) a herbivore that in turn is consumed by (iii) a predator. Our results highlight that predation may result in local accumulation of the resource via top-down control of the herbivore. As such, the temporal and spatial variation of the resource distribution is increased, selecting for increased herbivore movement. This results in selection of larger herbivores than in the scenario without predator. As predators cause herbivores to be intrinsically much larger than the optimal sizes selected by habitat fragmentation in the absence of predators, habitat fragmentation is no longer a driver of herbivore size. However, there is selection for increased predator size with habitat fragmentation as herbivores become less abundant, favoring gap-crossing ability of the predator. Since herbivore and predator body size respond differently to habitat loss and fragmentation, realized predator-herbivore body size ratios increase along this fragmentation gradient. Our model predicts the dominance of top-down forces in regulating body size selection in food webs and helps to understand how habitat destruction and fragmentation affect overall food web structure.

## Introduction

Body size represents a super trait, regulating almost any trait of an individual by its effect on metabolic rate (Peters, 1983; Brown *et al.*, 2004; Fritschie and Olden, 2016; Brose *et al.*, 2017). As such, an individual’s behavior, ecology and function are constrained by its body size (Bartholomew, 1982; Peters, 1983; Brown *et al.*, 2004). For example, small individuals have short generation times and low energetic requirements whereas large individuals have higher average speed of movement and resource consumption (Peters, 1983; Hirt *et al.*, 2017).

Herbivore species can show different allocation strategies: either few large or many small herbivore individuals can exist given a certain amount of resources (Delong and Vasseur, 2012; Yeakel, Kempes and Redner, 2018). This observation that the cost of total metabolic biomass is independent of body size is known as the ‘Energetic Equivalence Rule’ (Atkins et al., 2015; Delong and Vasseur, 2012; Yeakel, Kempes and Redner, 2018; Damuth, 1981). However, the total metabolic biomass of a herbivore species is constrained by resource availability or bottom-up dynamics. Importantly, with increasing trophic level, more complex size-dependent processes imply extra energetic and mechanical constraints. For a predator, prey that is too small are difficult to locate and render little energy, whereas prey that is too large might be hard to control and capture (Brose *et al.*, 2006; Portalier *et al.*, 2018). In foraging theory, this trade-off is represented by a hump-shaped function for predation rate, with a maximum at intermediate predator-prey ratios (Brose *et al.*, 2008). As such, predator-prey body size ratios are optimized in relation to habitat, prey and predator type, depending on the specific costs and constraints of the system (Brose *et al.*, 2006). Generally, these constraints and limits result in predators that are larger than their prey (Brose *et al.*, 2006; Portalier *et al.*, 2018), corresponding to one of the earliest observations in biology (Elton, 1927). By preferentially consuming prey of specific sizes, predators thus exert top-down forces within a food web (Howeth *et al.*, 2013). The emerging predator-prey body size ratios are theoretically demonstrated to maximize food web stability (Emmerson and Raffaelli, 2004; Brose, Williams and Martinez, 2006). In a tri-trophic food web (Otto et al., 2007), for instance, deviations from optimal predator-prey body sizes lead to predator extinction or unstable overshooting dynamics by resource accumulation (as described by the paradox of enrichment) (McCann, 2012). When predators are much smaller than their prey, energetic demand will increase during foraging (higher mass-specific metabolic rate with decreasing size), resulting in predator extinction by resource limitation (Otto et al., 2008). On the contrary, when predators are much larger than their prey, prey will eventually be suppressed, thereby giving rise to basal resource accumulation (Otto et al., 2008).

Because individual movement capacities and efficiencies are strongly related to energy use and body size, the spatial distribution of resources will impose selection on body size (Allen *et al.*, 2006; Hirt et al. 2018). Selection favors those individuals that move at a spatial scale at which resources are abundant and ensure optimal resource access (Holling, 1992; Nash *et al.*, 2014; Raffaelli *et al.*, 2016). Because of the current threat of habitat loss and fragmentation, many species are expected to experience changes in the spatial organization of their habitat, and these are thought to be at the basis of many observed body size shifts. Body size shifts due to habitat fragmentation are widely documented in nature but so far not well understood (Lomolino and Perault, 2007; Braschler and Baur, 2016; Renauld *et al.*, 2016; Warzecha *et al.*, 2016; Merckx *et al.*, 2018). Habitat loss refers to a decrease in the amount of suitable habitat whereas fragmentation per se implies a decrease in the spatial autocorrelation of suitable habitat (Jackson and Fahrig, 2013; Fahrig, 2017). It is important to study both effects independently, as each has a distinct effect on species performance within multi-trophic food webs (Liao, Bearup and Blasius, 2017). Habitat loss generally has negative effects on species survival, whereas fragmentation might promote species coexistence within a trophic level by lowering competition and between trophic levels by providing refuges (Jackson and Fahrig, 2013, 2015; Fahrig, 2017; Liao, Bearup and Blasius, 2017; Fletcher Jr *et al.*, 2018). So far, theoretical studies have demonstrated that large individuals can be selected with increasing levels of isolation and habitat fragmentation due to their high gap-crossing ability (Etienne and Olff, 2004; Hillaert, Hovestadt, *et al.*, 2018). Within a resource-consumer context, however, this selection of large individuals has only been observed in case of completely informed movement that lowers the risk of arriving in unsuitable habitat (Hillaert, Vandegehuchte, *et al.*, 2018). Whether predators respond similarly to habitat loss and fragmentation as their prey is unclear. While the resource of the prey is stationary, the resource of the predator is mobile; selection on herbivore and predator size during habitat loss and fragmentation may thus be different. Moreover, predators exert strong selection on consumer body size by consuming only particular sizes according to their preferred optimal predator-prey body size ratio (Howeth *et al.*, 2013; Tsai, Hsieh and Nakazawa, 2016). In order to consider the top-down effect of the predator on consumer selection, it is essential to include food web topology in studies of species responses to habitat fragmentation (Liao, Bearup and Blasius, 2017). Importantly, differential body size responses across trophic levels might shift realized predator-prey body size ratios (Tsai, Hsieh and Nakazawa, 2016), thus affecting predator-prey interaction strength (Emmerson and Raffaelli, 2004).

As mentioned before, theory so far focused on how body mass distribution shifts within one trophic level (Milne *et al.*, 1992; Etienne and Olff, 2004; Buchmann *et al.*, 2011, 2013; Hillaert, Hovestadt, *et al.*, 2018). However, this study does not include the effect of predation. To increase realism, we here studied the effect of habitat loss and fragmentation on body size selection within a simple resource-herbivore-predator model. This was achieved by extending the model presented in (Hillaert, Vandegehuchte, *et al.*, 2018) with an extra trophic level. In this model, individual traits of the herbivore and predator are described by established allometric rules (Peters, 1983). We focus on the effect of habitat fragmentation at the scale of foraging while distinguishing the process of habitat loss from the process of fragmentation per se. The scale of foraging is applied because fine-grained fragmentation has a larger effect on individual survival and reproduction than coarse-grained fragmentation when a species invests more time in foraging than dispersing (Cattarino, Mcalpine and Rhodes, 2016). Our goal is to answer the following questions: (i) How does predation affect body size selection in the herbivore? (ii) Do trophic levels respond differently to fine-scale habitat fragmentation and destruction? (iii) Which effects dominate: top-down or bottom-up? (iv) Are realized predator-prey body size ratios affected by fine-scale habitat fragmentation and destruction?

## Material and methods

We here took an arthropod-centered approach and parameterized allometric rules for a herbivore and predator that are both haploid and parthenogenetic with a semelparous lifecycle.

By applying an individual-based approach, we were able to include intra-specific size variation and stochasticity within our model. This approach in conjunction with the assumption of asexual reproduction and equivalent ontogenetic and interspecific scaling exponents (West, Brown and Enquist, 2001; Moses *et al.*, 2008), implies that our results can be interpreted both at the metapopulation and metacommunity level for both the herbivore and the predator. A detailed description of the model following the ODD (Overview, Design concepts and Details) protocol is available in supplementary material part 1 (Grimm *et al.*, 2010). The applied model is based on Hillaert, Vandegehuchte, *et al.* (2018).

### The landscape

The landscape is a cellular grid of 200 by 200 cells and is generated using the Python package NLMpy (Etherington, Holland and O’Sullivan, 2015). Each cell within the landscape has a side length (*SL*) of 0.25 m and therefore a total surface of 0.0625 m^2^. Within the landscape, a distinction is made between suitable and unsuitable habitat. Only within suitable habitat, the basal resource is able to grow. When testing the effect of landscape configuration, the proportion of suitable habitat (*P*) and habitat autocorrelation (*H*) were varied between landscapes. Habitat availability increases with *P*, whereas habitat fragmentation decreases with *H.* The following values were assigned to *P*: 0.05, 0.20, 0.50 or 0.90. *H* equaled either 1 (in all four cases), 0.5 (when *P* equaled 0.05 or 0.20) or 0 (when *P* equaled 0.05). As such, highly fragmented landscapes with a high amount of suitable habitat were not included in the analysis as these rarely occur in nature (Neel, McGarigal and Cushman, 2004).

### The basal resource

Local resource biomass is represented as the total energetic content of resource tissue within that cell (*R_x,y_* in Joule). This resource availability grows logistically in time depending on the resource’s carrying capacity (*K*) and intrinsic growth rate (*r*). In any cell, a fixed amount of resource tissue (*E_nc_*, in Joules, fixed at *2 J*) is non-consumable by the herbivore species, representing below-ground plant parts. As such, *E_nc_* is the minimum amount of resource tissue present within a suitable cell, even following local depletion by the herbivore species.

### Herbivore and predator

All herbivores and predators are modelled as individuals within the landscape. Both, herbivore and predator develop through two life stages: a juvenile and adult life stage. Within a day, both stages have the chance to execute different events (see Figure 1). Each day an individual executes all these events sequentially. The order in which individuals (herbivores and predators) are selected is randomized daily. Importantly, during the consumption event, the herbivore feeds on the basal resource whereas the predator feeds on the herbivore.

**Figure 1:**
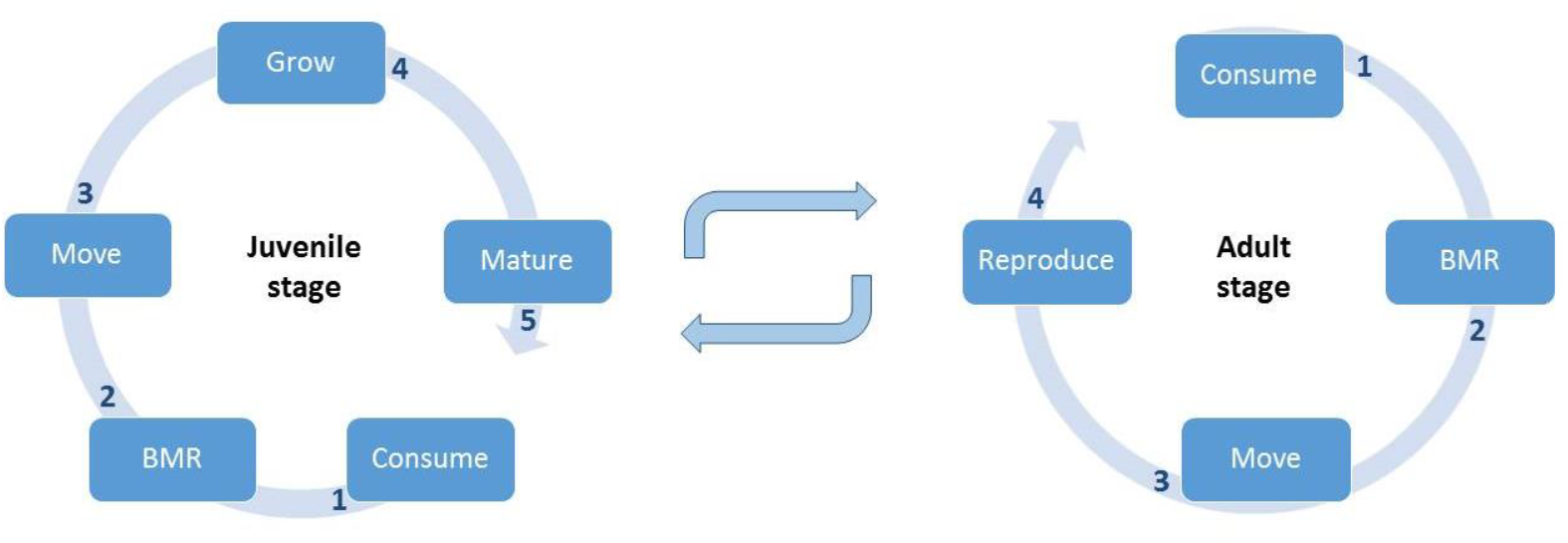
A comparison of daily events for the juvenile and adult stage of the herbivore and the predator (Hillaert, Vandegehuchte, *et al.*, 2018). BMR stands for basal metabolic rate costs. Numbers highlight the ordering of events within a day.

First, an individual nourishes its energy reserve by resource consumption and predating. Second, the energy reserve is depleted by the cost of daily maintenance (i.e. basal metabolic rate) and the cost of movement. Third, juveniles further deplete the energy reserve by growth, eventually resulting in maturation if they reach their adult size (*W_max_*). Energy that was not utilized is stored within the energy reserve. Adults can only reproduce if their internally stored energy (*E_r_*) exceeds a predefined amount. As the herbivore species and the predator species are semelparous, adults die after reproduction.

In both the herbivore and the predator, an individual’s body size at maturity (*W_max_*, in kg) is coded by a single gene. Adult size is heritable and may mutate with a probability of 0.001 during reproduction. A new mutation is drawn from the uniform distribution [*W_max_* – (*W_max_*/2), *W_max_* + (*W_max_*/2)] with *W_max_* referring to the adult size of the parent. New mutations may not exceed the predefined boundaries [0.01g, 3g] that represent absolute physiological limits. Both minimum and maximum weight are similar for the predator and the herbivore. New variants of this trait may also originate by immigration (see immigration below). Mutation enables fine-tuning of the optimal body size, whereas immigration facilitates fitness peak shifts.

### Initialization

For any parameter combination, 50 simulations were run. At the start of a simulation, adult individuals were introduced with an average density of one herbivore per two suitable cells. After 20 timesteps, 1000 predators are randomly added to the landscape in any suitable cell. This time lag allows the herbivore to reach a stable population size, increasing predator survival chances. The adult mass of each individual (*W_max_*) (for both herbivores and predators) was defined as ten raised to the power of a value drawn from the uniform interval [−5, −2.522878745]. In other words, we sample a value between 0.00001 kg (minimum adult mass) and 0.003 kg (maximum adult mass). As such, individuals with masses of different orders of magnitude have an equal chance of being initialized in the landscape. Moreover, initialized distributions are skewed to small individuals. Initial resource availability per cell was 100 J. Total runtime was 3000 time steps for all scenarios, with one time step corresponding to one day.

### Immigration

The frequency with which predator and herbivore immigrants arrive in the landscape is described by *q*. This variable is fixed at one per 10 days. The process of determining an immigrant’s adult mass is similar as during initialization (see above). An immigrant is always introduced within a suitable cell and its energy reserve contains just enough energy to cover the cost of basal metabolic rate and movement during the first day.

### The implementation of body size

The assumptions describing the daily events of the herbivore are described in the resource-consumer model (Hillaert, Vandegehuchte, *et al.*, 2018). Some events do not differ significantly between trophic levels and are therefore assumed to be identical for the herbivore and the predator (this is the case for basal metabolic rate, growth, maturation and reproduction. Details are provided in (Hillaert, Vandegehuchte, *et al.*, 2018) and the ODD protocol in supplementary material part 1. The events that differ between the predator and the herbivore are described below.

### Consumption

Individual ingestion rate (*IR*, in Watts) of an individual increases with its size (*W*, in kg) by the following equation for both the herbivore and the predator:

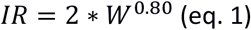

Following log transformation, the slope (0.80) was found by Peters (1983) to be the mean of several studies focusing on ingestion rates of poikilotherms (Peters, 1983). The intercept of this equation lays within the observed range of elevations [0.12 to 2] of these studies (Peters, 1983).

Based on eq. 1, the amount of energy ingested per day for an individual (*i_max_* in Joules) is determined as

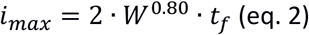

with *t_f_* referring to the time devoted per day to consumption (in seconds), which is fixed at 15 hours.

#### The herbivore

The amount of resources consumed by a herbivore (*E_c_*) only equals *i*_max_ if this amount is available. Otherwise, *E_c_* equals the amount present within a cell. As such, we assume contest competition for resources, with a competitive advantage for those individuals which are randomly selected first during a day.

When we consider that the herbivore feeds on young terrestrial foliage, it can only assimilate 65 percent of its daily ingested energy (Ricklefs, 1974 cited in Peters, 1983). Moreover, we assume that the herbivore loses 10 percent of its ingested energy to processing costs (i.e. specific dynamic action) (Ricklefs, 1974). As such, only 55 percent of the ingested energy remains available to the organism. Therefore, the energy that is being assimilated by a herbivore individual (*E_a_* in Joules) is described by

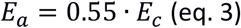

#### The predator

For each predator, the herbivore individuals located within its cell are selected within a random order. Per selected herbivore, the chance of successful attack (*s_a_*) is calculated. This chance is defined by multiplying the chance of interaction based on herbivore abundance (*i_PH_*) with a measure for optimality of the predator-herbivore body size ratio (*O_BSR_*):

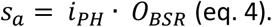

*i_PH_* increases with herbivore abundance in a cell, according to:

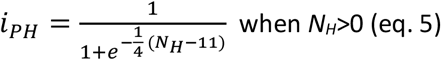

with *N_H_* representing the number of herbivores present within a cell, being continuously updated during a day. This function has a sigmoid shape and therefore implies a functional type III response (see Figure 2), stabilizing food web dynamics as highlighted by the sensitivity analysis (see supplementary material part 2). During a day, the number of herbivores present in a cell (*N_H_*) is constantly updated.

**Figure 2:**
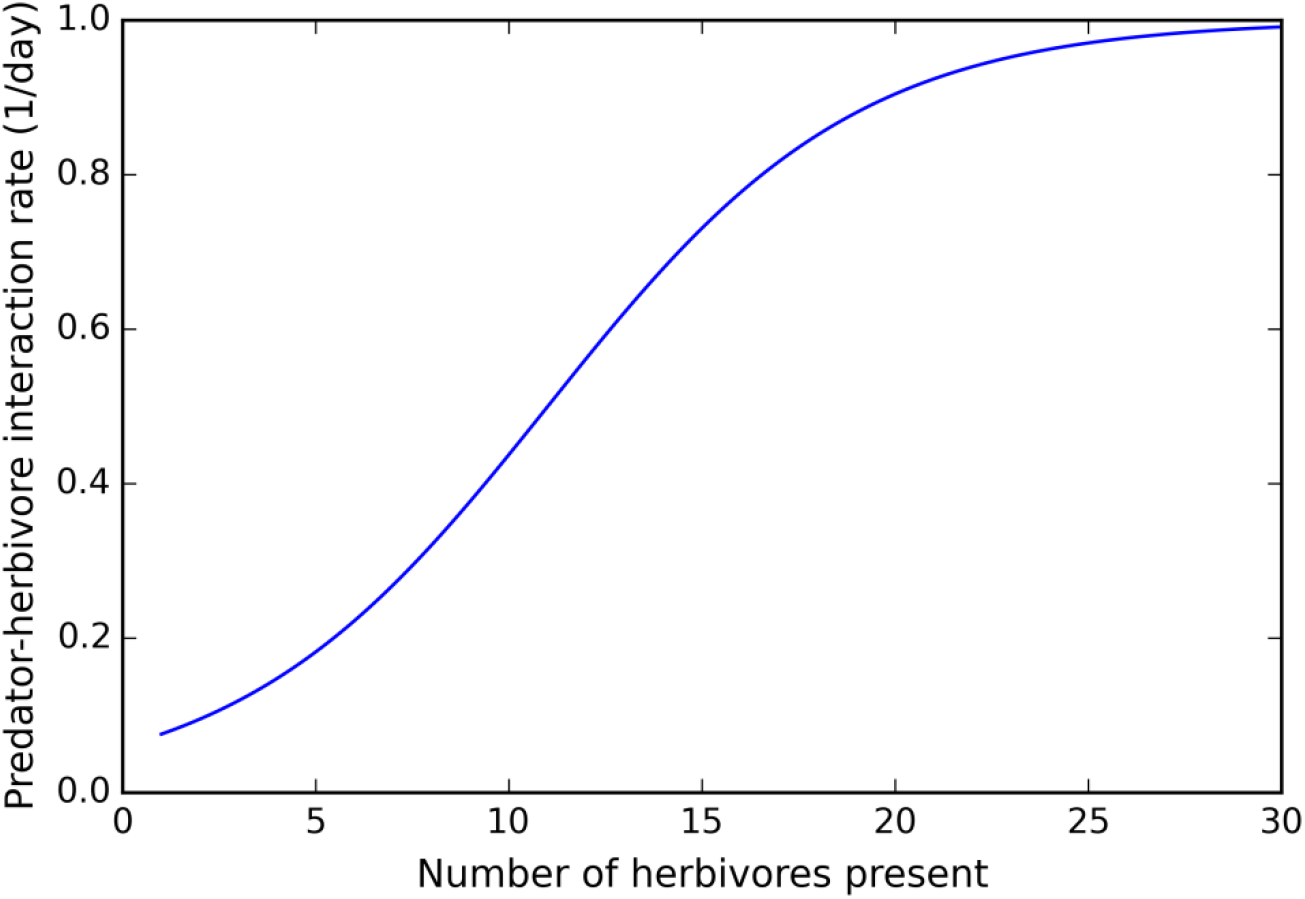
Relationship between predator-herbivore interaction rate and number of herbivores present within a cell. During a day, the number of herbivores present in a cell is constantly updated.

Contrary to a preferred predator-prey body mass ratio which depends on predator body mass, we included a fixed ratio which is in line with (Tsai, Hsieh and Nakazawa, 2016). Per selected predator-herbivore pair, the corresponding log_10_(predator-herbivore body mass ratio) is calculated. This ratio is then compared with the observed distribution of log_10_(predator-prey body mass ratios) in terrestrial systems with invertebrate predators (normal distribution with average 0.6 and SD 1.066) (Brose *et al.*, 2006). We refer to this observed distribution as the preferred predator-herbivore body mass ratio (Tsai et al., 2016). If the ratio of the selected pair is rarely observed in nature, the value for *OBSR* is close to zero. In case the ratio is often observed, the value for *OBSR* lays close to 1. In order to obtain values for *O_BSR_* between 0 and 1, the observed normal distribution in nature is scaled by an extra factor. As such, the formula for the calculation of *O_BSR_* is the following (see Figure 3):

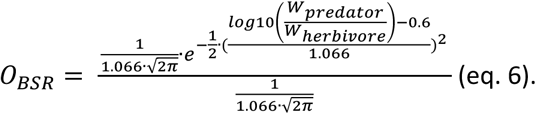

**Figure 3:**
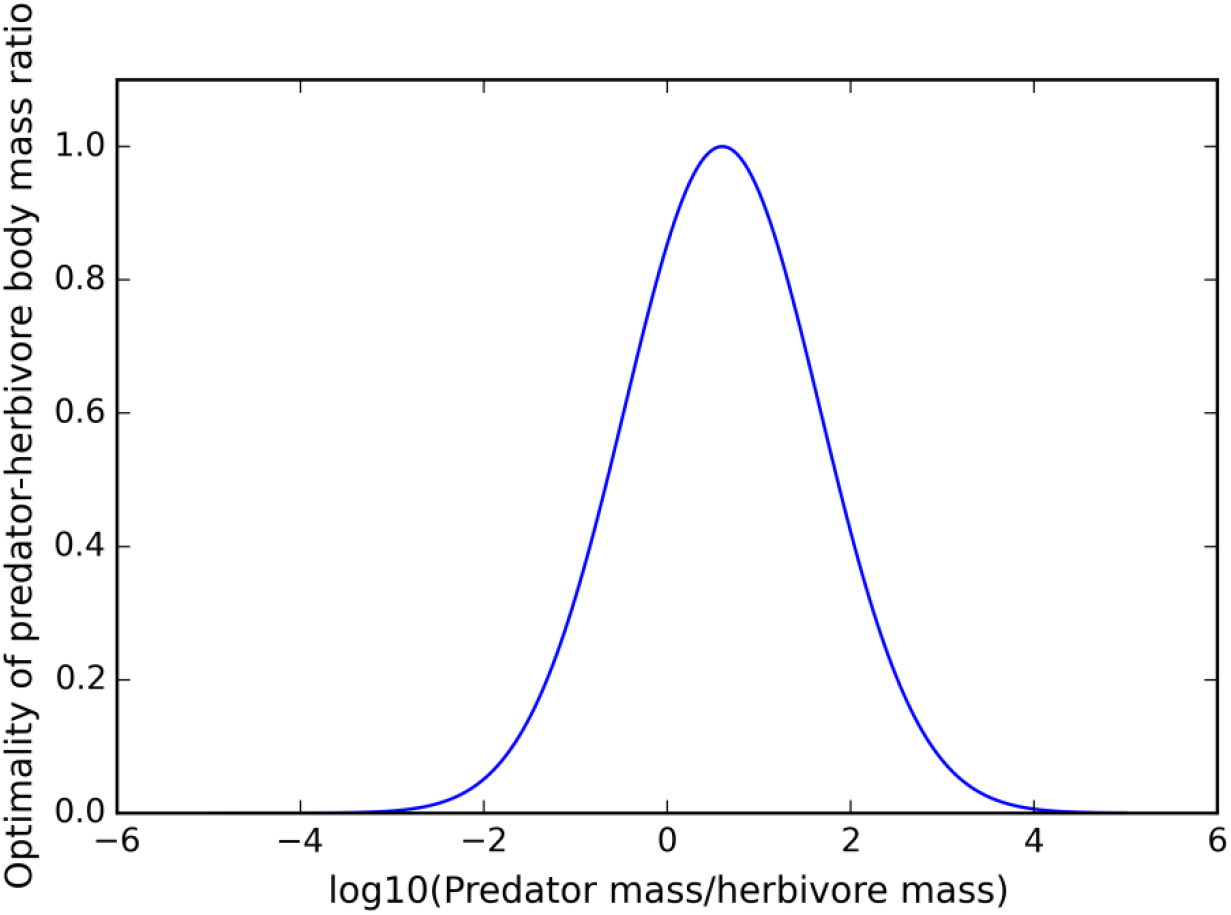
The implemented optimal predator-herbivore body size ratio is displayed. This distribution is observed by Brose (2006) to occur in terrestrial systems for invertebrate predators.

As *i_PH_* and *O_BSR_* are both numbers within the interval [0,1], the same is true for *s_a_*. In case a randomly sampled number from the interval [0,1] is smaller than *s_a_*, the attack of the predator on the herbivore is successful and *E_c_* of the predator is increased with *W_t,herbivore_* *7000000 + *E_r,_*_herbivore_. This formula assumes that the energetic content of wet tissue corresponds to 7 × 10^6^ Joule per kg (Peters, 1983) and that the body mass of a herbivore (*W_t,herbivore_*) does not include the energy stored within its energy reserve (*E_r_*). As long as *E_c_* is smaller than *i_max_* of the predator, another herbivore within the same cell may be attacked by the predator. However, *E_c_* does never exceed *i_max_*.

Considering that the predator feeds on insects, it may assimilate 80 percent of its daily ingested energy (Ricklefs, 1974; Peters, 1983). However, we assume that the predator loses 25 percent of its ingested energy to processing costs (i.e. specific dynamic action) (Ricklefs 1974 cited in Peters 1983). As such, only 55 percent of the ingested energy remains available to the organism. Therefore, the energy that is being assimilated by a predator individual (*E_a_* in Joules) is described by the same formula as for the herbivore (see eq. 3).

## The movement phase

### Probability of moving (*p*)

Whether an individual moves, depends on the ratio of the amount of energy present within a cell relative to the amount of energy it can eat during a day (*i_max_*).

The probability of moving (*p*) for a herbivore is thereby calculated as, based on Poethke and Hovestadt, 2002:

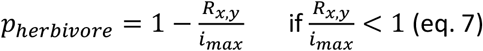

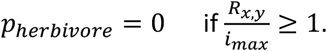

A predator’s probability of moving is based on *s_a_*: the chance of moving decreases with the chance of successful attack by

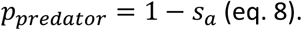

In the formula of *s_a_*, the average herbivore mass within the cell is applied (see eq. 4 and 6).

### Defining searching area

As one time step in our model corresponds to one day, we do not model the movement behavior of an individual explicitly, but instead estimate the total area an individual can search for resources during a day. This area is called an individual’s searching area is calculated once per time step, for each moving individual. As all cells within a particular distance from the origin are equally intensively searched, the searching area is circular with a radius (*rad*) and a center corresponding to the current location of an individual (Delgado *et al.*, 2014). An individual’s searching area increases with an individual’s optimal speed (*v_opt_*), movement time (*t_m_*) and perceptual range (*d_per_*). Both optimal speed and perceptual range depend on body mass, resulting in larger searching areas for larger individuals. The cost of movement includes the energy invested by an individual in prospecting its total searching area. Therefore, it is dependent on the size of the total searching area instead of the shortest distance between the cell of origin and cell of destination.

An individual’s optimal speed of movement (*v_opt_*, in meters per second) is calculated for herbivores according to the following equation, derived for walking insects (Buddenbrock, 1934; Peters, 1983):

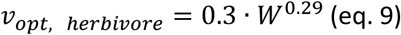

Speed of movement (*v_opt_*, in meters per second) of the predator is defined by the following equation (Hirt *et al.*, 2017):

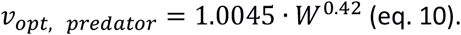

The time an individual invests in movement per day (*t_m_*, in seconds) is maximally 1 hour. In case too little internally stored energy is present to support movement for one hour, *t_m_* is calculated by:

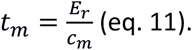

*c_m_* refers to the energetic cost of movement (in joules per second) and is calculated for herbivores by the following formula, which is based on running poikilotherms (Buddenbrock, 1934; Peters, 1983)):

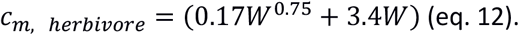

We adapt the formula of *c_m_* for the predator by implementing the formula for *v_opt_, _predator_* in the formula of *c_m_* (see supplementary material part 3 for derivation):

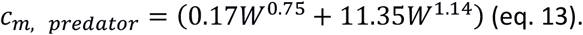

The cost of moving during the time *t_m_* (*c_m_⋅t_m_*) is subtracted from an individual’s energy reserve. Based on *t_m_* and *v_opt_*, the total distance an individual covers at day *t* (*d_max_*) is determined as:

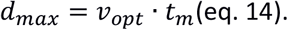

Next, the perceptual range of an individual is determined by means of the following relationship:

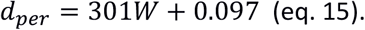

For simplicity, this relationship is linear and based on the assumption that the smallest individual (0.01g) has a perceptual range of 0.10 m and the largest individual (3g) a perceptual range of 1m. The effect of this relationship has been tested (see supplementary material part 2). Moreover, the positive relationship between body size and perceptual range or reaction distance has been illustrated over a wide range of taxa, including arthropods (supplementary information of Pawar, Dell and Van M. Savage, 2012).

The foraging area of an individual is circular and its radius (*rad*, in m) is calculated by taking into account the total distance the individual has covered during the day and the individual’s perceptual range (see Supplementary material part 5 for explanation of this formula):

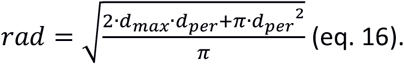

In order to avoid side-effects of applying the variable rad for a continuous landscape within a cellular landscape, a randomly drawn value from the following distribution, [−0.5 ∙*SL*, 0.5 ∙ *SL*], is added to *rad*.

### Habitat choice

Here, movement is informed as an individual always moves to the cell with the highest amount of resources (the herbivore) or the cell with the highest rate of successful attack (based on average herbivore weight per cell in case of the predator) within its foraging area.

### Output

Only simulations in which the predator persists during the final 500 days of a simulation are included in the analysis. An overview of the number of included simulations per landscape type is given in Table S2.1. During each simulation, we traced changes in the mean amount of resources per cell and total number of adults and juveniles and average adult mass (*W_max_*) of both the herbivore and the predator over time. Throughout the final 1500 days of a simulation, 1000 eggs (for predators and herbivore each) were randomly selected to be followed during their lifetime. The movements and reproductive success of the resulting herbivore individuals were recorded. During the final 100 days of a simulation, the log_10_(predator-herbivore body mass ratio) was recorded per successful predation event. As such, the average log_10_(predator-herbivore body mass ratio) could be determined per scenario, as well as the deviation of this average from the implemented optimum log_10_(body mass ratio).

At the end of a simulation, the body masses of maximally 50 000 predators and maximally 50 000 herbivores were randomly sampled. Also, the abundance of predators and herbivores as well as the resource amount per cell was written out. This enables us to study the spatial distribution of the predator(s), the herbivore(s) and the resource.

In order to determine the effect of the predator(s) on herbivore body weight distributions, the settings of the resource-herbivore-predator model were applied to run a comparable model without predator (see Table S1.1).

## Results

In each landscape type, the body mass of the predator is selected to be higher than that of the herbivore. Habitat loss, in conjunction with fragmentation, selects for an increase in average body mass of the predator (Figure 4). Habitat loss within highly autocorrelated landscapes (*H* equaling 1), does not clearly affect average predator body mass. However, the number of simulations in which the predator and herbivore survive during the final 500 days of a simulation are lowest when *P* equals 0.05 and *H* 1 or *H* 0.05 (Table S2.1). Although a similar pattern is observed for herbivore body mass when no predator is present, herbivore body mass shows almost no response to habitat fragmentation in the presence of a predator. This pattern is always supported by the sensitivity analysis, except for the scenario with a clutch size of 2. Furthermore, in case of *P* 0.05 and *H* 1, average predator body mass sometimes approaches that of the scenario with *P* 0.05 and *H* 0. When this is the case, the number of included simulations is low due to extinction of the predator. Moreover, in this landscape type (*P* 0.05 and *H* 1), drift is strong, explaining the variation in average body mass between simulations. Notably, the body mass of a herbivore is overall larger when a predator population or community is present, except for the landscape with *P* equaling 0.05 and *H* 0.

**Figure 4:**
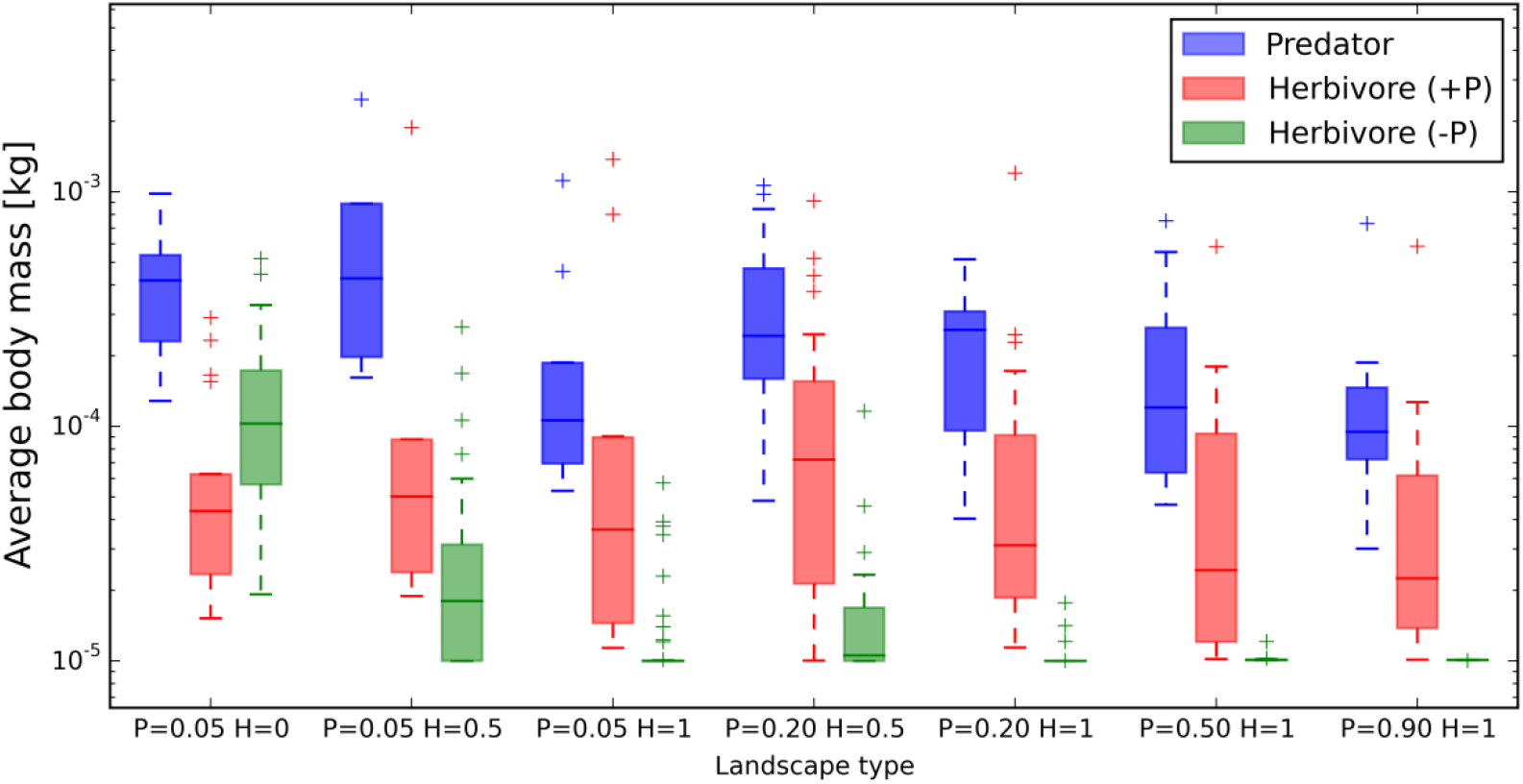
The effect of habitat loss and fragmentation of a resource on average body mass of its herbivore and a predator. In order to infer the effect of predation, average herbivore body mass is also displayed for a scenario in which the predator was not present (see legend). For an overview of the number of simulations per scenario, see Table S2.1 in supplementary material part 2. An overview of the parameter settings is given in Table S1.1 in supplementary material part 1.

Temporal and spatial dynamics of the resource and the herbivore are strongly affected by the presence of a predator, illustrating the strength of the top-down force. Dynamics within the predator-herbivore-resource food web fluctuate strongly over time (Fig S4.1). Moreover, the spatial distribution of the resource and the herbivore is highly heterogeneous (Fig 5). When a predator is present, the number of suitable patches occupied by the herbivore is lower (Fig S4.2). Also, the average amount of resources per cell is higher (Fig S4.3), and even local accumulation occurs (Fig 5). Importantly, top-down and bottom-up forces strongly interact in our model. For example, resources increase in abundance with habitat fragmentation and destruction when a predator is not present (Fig S4.3). In contrast, habitat fragmentation and destruction result in a decrease in resource amount when a predator is present (Fig S4.3).

**Figure 5:**
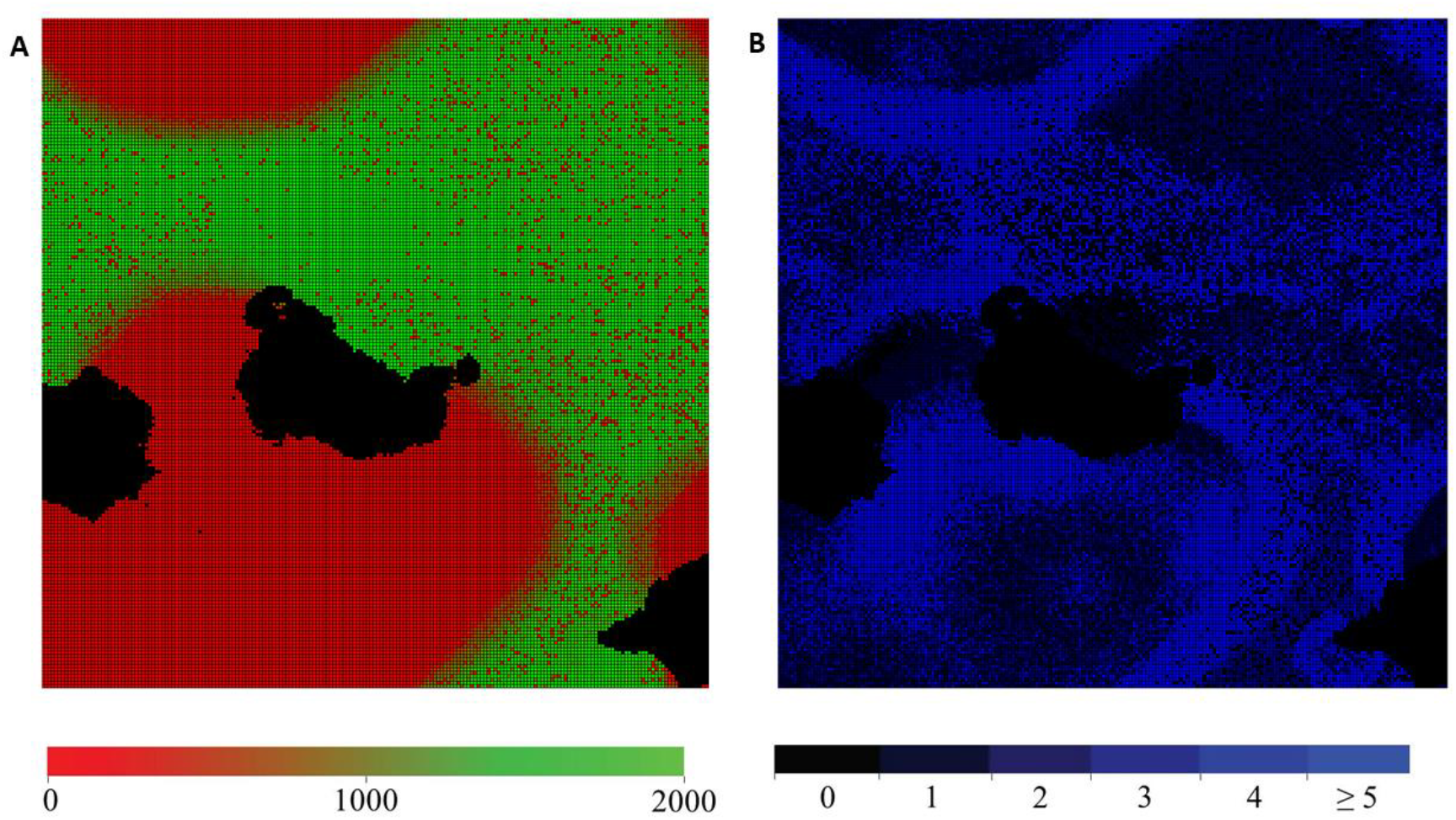
Spatial distribution of the herbivore (number of individuals) and resource (in Joule) within one simulation with P equaling 0.90 and H 1 when a predator is present. A) The distribution of resources is displayed (black: unsuitable habitat) and B) the distribution of herbivores (black: no herbivores present).

The average realized log_10_(predator-herbivore body mass ratio) strongly approximates the preferred ratio when *P* equals 0.9 and *H* equals 1 (Figure 6). However, with increasing habitat loss and fragmentation, the realized log_10_ (predator-herbivore body mass ratio) is selected to increase, up to a maximum at *P* = 0.05 and *H* = 0 (Figure 6). This deviation from the preferred ratio with increasing habitat loss and fragmentation is strongly confirmed by the sensitivity analysis (Table S4.1). Moreover, the sensitivity analysis highlights that parameter changes that limit movement increase the overall deviation (Table S4.1), while parameter changes that facilitate movement decrease the deviation (e.g. higher value for *t_m_*) (Table S4.1).

**Figure 6:**
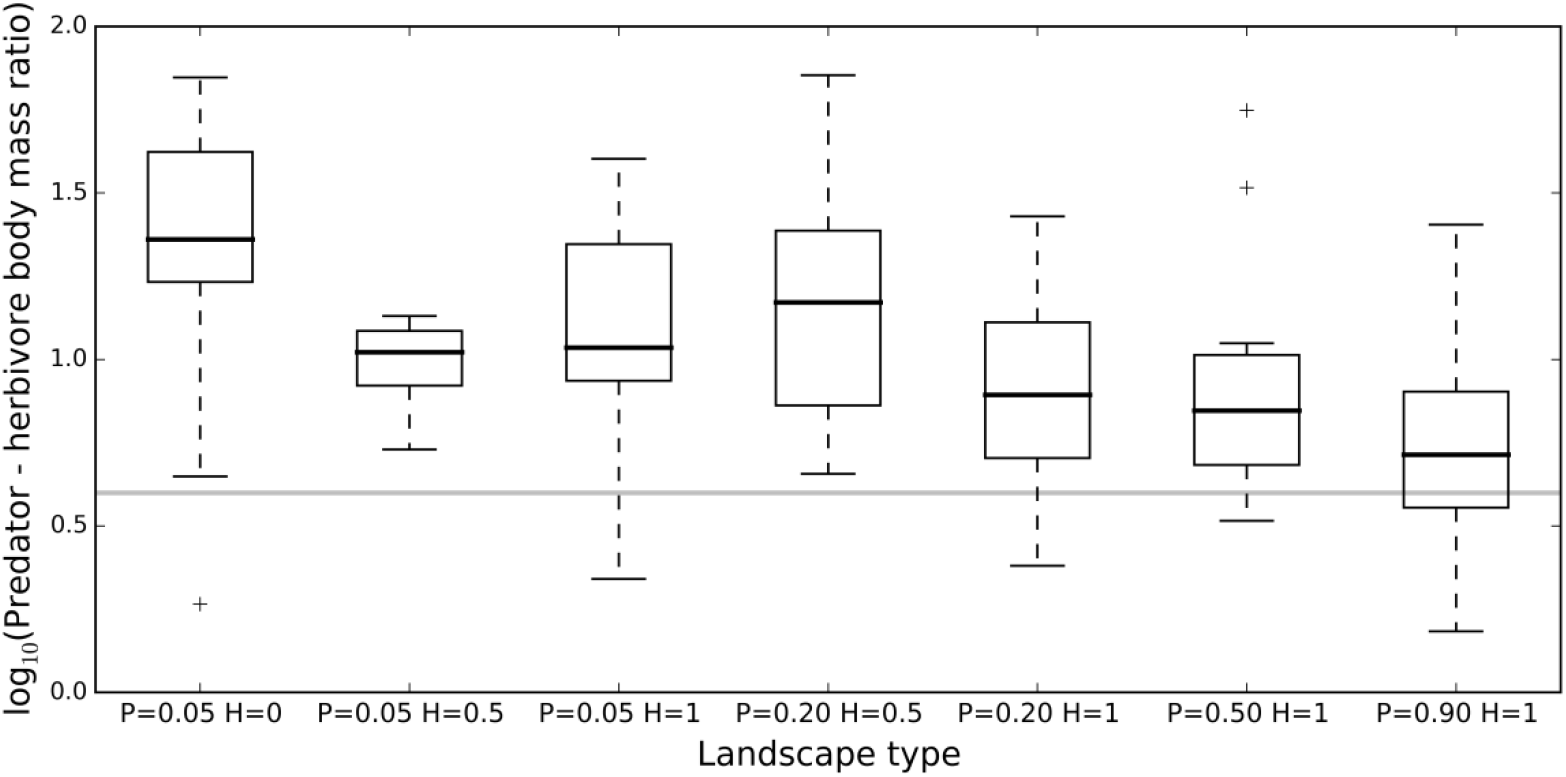
The effect of habitat loss and fragmentation on average realized predator-herbivore body mass ratios. The horizontal line represents the preferred predator-herbivore body mass ratio maximizing the predators’ foraging success. For an overview of the number of simulations per scenario, see Table S2.1 in supplementary material part 2. An overview of the parameter settings is given in Table S1.1 in supplementary material part 1.

## Discussion

First principles from movement ecology and metabolic theory predict how fine-grained habitat fragmentation changes selection on body size within a simple three-trophic food web model. The findings of our model are the following. (i) Predators induce a spatially and temporally heterogeneous distribution of the resource, thereby selecting for increased movement (ability) and thus increased size in herbivores. (ii) Predators cause herbivores to be intrinsically much larger than the optimal sizes selected by habitat fragmentation in the absence of predators, so that habitat fragmentation is no longer a driver of herbivore size. Since habitat fragmentation causes herbivore abundance to decrease, it selects for a large predator size as larger predators are more mobile. (iii) Body size distributions of primary consumers are largely regulated by top-down forces. (iv) The realized predator-prey body size ratio increases with habitat fragmentation due to different selection at different trophic levels.

### Effect of predators on herbivore size

In the absence of predators, selection on herbivore body size has been demonstrated to depend on the spatial organization of resources, and information use during movement (Hillaert, Vandegehuchte, *et al.*, 2018). Without predator interactions, the optimal body mass of herbivores that move in an informed way increases with habitat fragmentation and loss. Moreover, when the percentage of suitable habitat is high, a herbivore’s body mass is minimized. Under these conditions, small herbivores are selected as these have the shortest generation times whereas no benefit results from being able to cover a large spatial extent and, hence, from being large, as resources are uniformly distributed in space. We here show that if a herbivore coexists with its predator, the herbivore’s temporal and spatial dynamics are much more unstable and resources become highly heterogeneously distributed in space. This arises because predators can deplete local herbivore populations, thereby enabling resource accumulation and generating high spatial and temporal variability in resource levels. As such, selection acts in favor of those herbivores that can reach cells with high amounts of resources first (Hastings, 1983). Hence, herbivores which move in an informed way are selected to be larger in the presence than in the absence of a predator. Since Amarasekare (2016) retrieved similar adaptive dynamics for dispersal in a simple tri-trophic foodweb, we can conclude that, here, selection for enhanced movement is the main driver behind body size evolution.

### Effect of habitat fragmentation on body size across trophic levels

In the absence of predators, herbivore size is selected to increase with habitat loss and fragmentation (Hillaert, Vandegehuchte, *et al.*, 2018). This effect disappears in our tri-trophic model, in the presence of predator-prey dynamics. Absence of a selection differential implies the presence of a single optimal herbivore size irrespective of the resource’s spatial organisation. As such, herbivores shift towards larger sizes in the presence than in the absence of predators when resources are abundant, but to smaller sizes when resources are rare and highly fragmented (*P* 0.05 and *H* 0). This inverse pattern can be explained by fitness disadvantages for the herbivore of being too large, associated with an increased time until maturity and hence increased lifetime predation pressure.

In contrast to the herbivore, the predator is always selected to be larger than the herbivore and, more importantly, its average body size increases with habitat fragmentation. The model observation that predators are larger than their prey follows logically from the implemented optimal predator-herbivore body mass ratio as observed in nature. Too high or too low predator-prey body mass ratios are not favorable as too small prey are hard to trace and offer low energy profit, whereas too large prey may be hard to control and capture (Brose *et al.*, 2006; Brose, 2010; Portalier *et al.*, 2018). Moreover, as predators need to keep track of mobile herbivores, selection on movement should always be strong in active hunters. This is supported by our modeling results, as optimal predator sizes are always a little larger than expected, based solely on the implemented preferred predator-prey body size ratio (Figure 5). Since selection for mobility in the predator is largest in the most resource-deprived and fragmented landscapes where herbivore abundances are lowest, the largest predators are selected here. This pattern is general under a wide range of boundary conditions (see sensitivity analysis) except for the scenario in which clutch size for the herbivore and predator is low. When clutch size of the herbivore is low, the predator size is selected to be large when habitat is abundant (*P* equaling 0.90 and *H* 1) relative to when it is rare (P equaling 0.05 and *H* 1). By constraining clutch size, the growth speed of the herbivore population is lowered. As such, the herbivore population growth rates are reduced, promoting predator mobility even when *P* is high. This mechanism is confirmed by the observation that lowering resource growth speed within the resource-herbivore model also resulted in selection of larger herbivores (Hillaert, Hovestadt, *et al.*, 2018). Under low *P* and low herbivore reproductive values, the largest predators can no longer persist due to food limitation and selection turns towards smaller average predator sizes.

Our theoretical predictions are confirmed by some but not all experimental studies. For instance, within a fine-grained fragmentation study, the density of the largest species of ground beetles responded positively to fragmentation (Braschler and Baur, 2016). However, in other predatory invertebrate species (spiders and rove beetles), response to fine-scale fragmentation was unrelated to body size (Braschler and Baur, 2016). In another study, web spiders showed no response to urbanization, which is associated with habitat fragmentation, whereas the community-weighted average body size decreased with urbanization in ground beetles and ground spiders (Merckx *et al.*, 2018). These and other counterintuitive outcomes might be explained by confounding factors. For instance, fragmentation due to urbanization is also linked with increasing temperatures by urban warming (Merckx *et al.*, 2018). Further, body size responses to habitat fragmentation might strongly be influenced by food web structure or the level of informed movement (Liao, Bearup and Blasius, 2017; Hillaert, Vandegehuchte, *et al.*, 2018). Generally, more experimental research on the effect of fine-scale fragmentation on body size across trophic levels is necessary to validate theoretical expectations, for instance by using the Metatron platform (i.e. an innovative infrastructure to study terrestrial organism movement under semi-natural conditions, Legrand *et al.*, 2012).

### Top-down versus bottom-up effects

Temporal and spatial dynamics of the resource and the herbivore are strongly influenced by the predator. The predator clearly suppresses herbivore population sizes at local scales and this effect cascades down the food web, resulting in a weaker control of the resource, which then locally accumulates. At this point, the top-down force influences the bottom-up one by creating temporal variation in resource abundance which imposes selection for larger and more mobile herbivores. This insight provides an explanation of why in a recent meta-analysis, top-down forces were found to be stronger than bottom-up forces for the fitness of terrestrial insect herbivores, considering that body size largely influences the fitness of an individual (Vidal & Murphy, 2018; Peters, 1983). However, the effect of bottom-up forces should not be underestimated. As highlighted by our modelling approach, habitat loss and fragmentation results in a selection for larger predator individuals whereas herbivore size does not respond. Consequently, predators are forced to consume herbivores that deviate from their preferred optimal size. Furthermore, we should note that movement of herbivores in our model is only influenced by the basal resource and not the predator, so non-lethal effects acting in landscapes of fear are not considered (Bleicher, 2017; Schmitz *et al.*, 2017). Moreover, we show that top-down and bottom-up act in concert and strongly interact. Without predators, habitat fragmentation prevents the consumer from reaching an ideal free distribution, hence imposing spatial variation in resource biomass (Hillaert, Vandegehuchte, *et al.*, 2018). As such, resources biomass increases globally with habitat fragmentation and destruction when a predator is not present. In contrast, when a predator is present, habitat fragmentation creates predator-free refuges for the herbivore. This increases the percentage of cells being occupied by the herbivore, globally controlling resource production. As such, habitat fragmentation and destruction decrease resource amount in the presence of a predator.

### Effect of habitat fragmentation on predator-herbivore body size ratio

Predators experience one extra selection pressure that is not experienced by the herbivore: predators are selected to have a size that approximates the preferred ratio to maximize chance of successful attack. Under the continuous availability of resources, in landscapes of *P* = 0.9 and *H* = 1, the selected predator-herbivore body mass ratio approximates the preferred ratio. However, as only predator size increases with habitat fragmentation, the available body mass distribution of herbivores deviates from the preferred one when resources are spatially structured (Tsai, Hsieh and Nakazawa, 2016). The realized predator-herbivore body mass ratio thus increases with habitat loss and fragmentation. Hence, the realized predator-prey body mass ratios and coupled interaction strengths are altered (Emmerson and Raffaelli, 2004). This model prediction coincides with the finding that prey limitation determines variation in predator-prey body mass ratios between food webs (Costa-Pereira *et al.*, 2018). Further, selection pressures that enlarge differences between preferred and available body mass distributions for predators might increase extinction rates of species from higher trophic levels. Moreover, our sensitivity analysis indicates that when predators are intrinsically more mobile (e.g. high *t_m_*), their realized predator-prey body mass ratio deviate less from the preferred ratio in highly fragmented landscapes. Whereas, when predators are intrinsically less mobile (e.g. low *t_m_*), their realized predator-prey body mass ratio deviate even more from the preferred ratio in these landscapes.

The predicted deviation of the predator-prey body mass ratio from the implemented optimum does consequently not only depend on the level of habitat fragmentation but also on the limitation by resources and the species-specific mobility traits.

## Conclusion

Our developed modeling framework, which merges principles from movement ecology and metabolic theory, shows that the effects of habitat fragmentation and destruction on body size distributions within food webs is not obvious. Predation selects for increased herbivore size by generating spatial and temporal variation in the distribution of the resource, favoring herbivore movement. As top-down forces dominate, the effect of predation should always be considered when estimating the effect of habitat fragmentation on changing selection pressures in food webs (Liao, Bearup and Blasius, 2017). Since predation results in larger optimal herbivore sizes in all landscape types, herbivore size no longer increases with habitat fragmentation as observed in a simpler consumer-resource food web. However, habitat fragmentation leads to larger optimal predator sizes as herbivores become rarer, favoring gap-crossing abilities and hence, movement potential, of the predator. Therefore, even if a herbivore and its predator persist under conditions of fine-scale fragmentation, the realized predator-herbivore body mass ratios will be larger than in continuous habitats. These deviations in realized predator-prey body mass ratios affect interaction strength, which may cascade through the food web and alter the energy flow (Emmerson and Raffaelli, 2004).

## Acknowledgements

JH was supported by Research Foundation – Flanders (FWO). The computational resources (STEVIN Supercomputer Infrastructure) and services used in this work were kindly provided by Ghent University, the Flemish Supercomputer Center (VSC), the Hercules Foundation and the Flemish Government – department EWI.

